# Understanding Coral Health from Reactor Engineering Perspective: Multiphysics Modeling of Coral–Environment Interactions

**DOI:** 10.64898/2026.01.21.700759

**Authors:** Haoyu Zhuo, Flora Lanxin Xiao, Xiao Dong Chen, Jie Xiao

## Abstract

Coral, as a bioreactor, has to continuously interact with surrounding environment to maintain a healthy state. A multi-physics reaction engineering model has been developed to capture this interaction. The coral interior is modeled as interconnected reaction units respectively for photosynthesis, respiration, and calcification, whose reaction kinetics are influenced by environmental fluctuations. Coupling between coral and environment is realized by bi-directional mass transfer at the coral-seawater interface, with consideration of the unique flow fields induced by ciliary beating. By resorting to this comprehensive model, we discover that ciliary beating demonstrates distinctively different diurnal and nocturnal functions. During daytime, beating can help reduce photosynthetic oxygen accumulation to prevent hyperoxia-induced mortality, while enhancing carbon dioxide uptake efficiency to promote nutrient production. At night, however, beating promotes oxygen acquisition for adequate respiration, while expelling carbon dioxide to inhibit symbiotic destruction under acidic stress.

The model further enables mechanistic analysis of the detrimental impact of climate change on coral health, where the influences from two key factors (i.e., temperature and CO_2_ level) can be decoupled. It*’*s interesting to find out that the elevated temperature plays a dominant role during daytime, while at night the coral is dominantly influenced by rising CO_2_ level.

## 1. Introduction

Coral reefs constitute an important part of the global marine ecosystem. Despite occupying only 0.5% of the seabed, these underwater ecosystems serve as habitats for about 30% of fish species [1]. Moreover, their ecosystem services such as coastal protection and food supply are essential for the daily lives of millions of people worldwide [2]. However, the ecological contributions of coral reefs are declining as the prominent environmental changes are inducing widespread mortality events [3].

Coral reefs are primarily constructed by numerous individual coral polyps [4], which cover the coral skeleton in a glove-like manner [5], forming the tentacle shaped organisms in contact with seawater in Fig. 1a. The dinoflagellate algae, also known as zooxanthellae (the green sphere inside the polyps in Fig. 1a), reside in the tissue of coral polyps. The symbiotic relationship between polyp and algae is crucial for their survival [6]. The zooxanthellae supply self-produced photosynthetic products to their coral host. The corals then utilize those products to synthesize nutrients and produce calcium carbonate for growth. In return, corals provide photosynthetic materials and shelter for the zooxanthellae [7].

**Figure 1.**
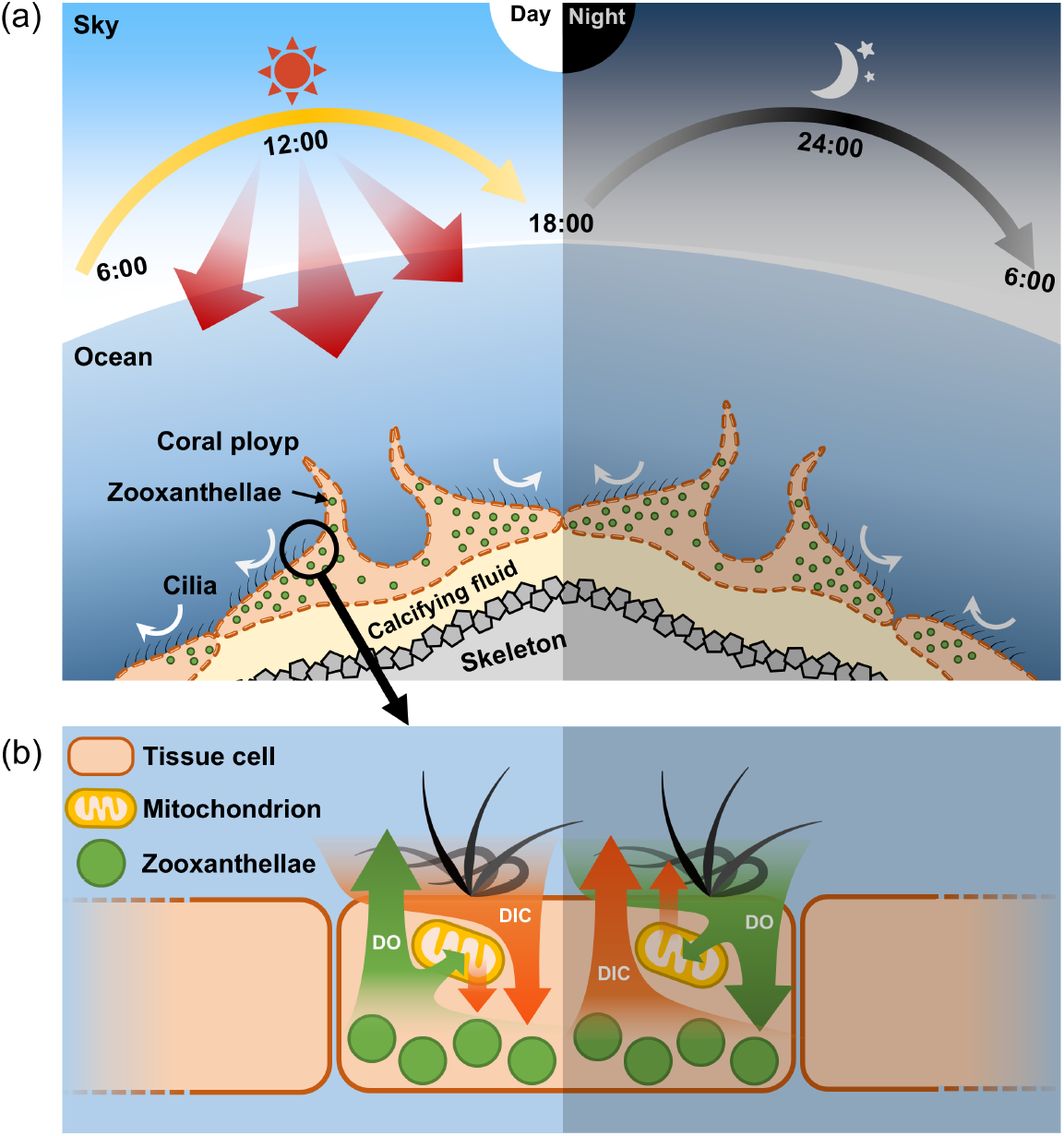
Schematic illustration of coral-environment interactions. (a) The complete system includes air, seawater, coral polyps covered with beating cilia with zooxanthellae inside the body tissue, calcifying fluid, and skeleton. Daytime configuration depicts sunlight exposure (left), while night-time configuration lacks solar irradiation (right). (b) Mass exchange between coral and environment. Photosynthesis and respiration take place inside the tissue cell. Ciliary beating drives localized fluid motion adjacent to the coral surface, enhancing mass exchange.

Although this win-win relationship sustained for millions of years, its stability is vulnerable to disruption, even minor environmental perturbations can destabilize this partnership [8]. When the symbiotic relationship is disrupted, colorful zooxanthellae are expelled from the coral tissue, leaving behind translucent tissues. This phenomenon, known as coral bleaching, manifests as a visible whitening of the calcium carbonate skeleton through the translucent tissues [9]. Global warming has been blamed to be a principal catalyst for this phenomenon, since the first documented global-scale bleaching event was in 1998, when a marked increase of seawater surface temperatures took place [10]. Additionally, other physical and biogeochemical conditions (e.g., light intensity, hypoxia, nitrogen enrichment, and bacterial infection) were also reported to compromise coral homeostasis, ultimately leading to bleaching [9].

It is known that coral bleaching is characterized by the destroy of the symbiotic relationship between zooxanthellae and their coral hosts. However, the complex relationships between the disruption of this symbiotic relationship and environmental changes remain unknown. Proposed hypotheses span from macro cellular [11], microbial [12], and smaller molecular scales [13].

From a chemical engineering perspective, corals are bioreactors composed of two interconnected sub-units, i.e., coral polyps and the zooxanthellae residing within their tissues. Physiological reactions occur within these biological structures, such as photosynthesis, respiration and calcification reactions, which synergistically determine the health status of both coral polyps and zooxanthellae. At the same time, the changing external environmental conditions, including temperature, atmospheric carbon dioxide levels, etc., affect the efficiency of these reactions. Furthermore, these living bioreactors engage in dynamic mass exchange with the surrounding seawater, thereby regulating their internal physicochemical conditions to maintain a suitable environment for survival. The exchange of dissolved oxygen (DO) and dissolved inorganic carbon (DIC) is shown in Fig. 1b. These mass exchanges, viewed as mass transfer processes from the chemical engineer perspective, are driven on one hand by concentration gradients between internal tissues and the external environment, and on the other by surface cilia, whose beating disturbs localized flow field next to the coral surface. In summary, to pursue in-depth and scientific understanding of coral health, coral and environment have to be considered as a holistic system.

Extensive experimental studies have been reported on the impacts of various environmental conditions on coral health. Notably, these environmental parameters include temperature [14], concentration of DO [15-17], partial pressure of atmosphere CO_2_ (pCO_2_) [18], light intensity [19, 20], seawater pH [21, 22], and even hydrodynamic conditions [23]. To quantitatively capture their influences, advanced monitoring techniques have been developed to trace both the physiological states of corals and the properties of the adjacent seawater. The specific parameters monitored include the spatial and temporal distributions of oxygen [24], pH [25], carbonate ion [26], as well as the spectral light field and light penetration in coral tissues [27]. However, due to the wide variety of the coral species and environmental conditions, the conclusions got from one set of experiments may not be applicable to other cases. Furthermore, decoupling the effects of multiple parameters in field experiments is almost impossible.

Mathematical models have also been developed to explore the ecosystems of coral reef. Specifically, coral-zooxanthellae-seawater coupled models have been established to depict both the symbiotic relationship of corals and zooxanthellae, as well as the mass exchange between coral and the surrounding seawater (28, 29). Wang et al. proposed an ecosystem dynamics model for predicting the impact of human activities on coral survival across several decades (30). However, those mathematical models are lumped ones that treat coral and its surrounding environment as a homogeneous system. Consequently, they are unable to capture the crucial spatial distributions that are especially important for scientific understanding of coral-environment interactions. Two-dimensional (2-D) or three-dimensional (3-D) CFD simulations were also developed to capture spatial distribution data. Wangpraseurt et al. simulated the oxygen distribution in seawater at the coral-algal interface by applying different surface photosynthesis and respiration rates (i.e., oxygen release flux) to the coral-seawater and algae-seawater interfaces, respectively [31]. In 2014, Shapiro et al. numerically investigated the regulatory effect of cilia beating on oxygen distribution in the seawater next to the coral surface under the assumption of a constant oxygen concentration on coral surface [32], which we found not true by analyzing their experimental data since experimental results show that ciliary beating can reduce O_2_ concentration on coral surface. Six years later, Pacherres et al. reported that ciliary beating does not influence oxygen release rates from coral [33]. Subsequently, a two-layer model assuming constant oxygen flux was developed to characterize the influence of ciliary beating on oxygen distribution [34]. However, the simplified assumption of constant oxygen concentration or flux on coral surface is questionable. Furthermore, it is still unknown how cilia regulate the transport of other substances beyond oxygen in order to maintain a healthy state. To explore phenomena inside the coral, Parkins et al. constructed a multiphysics model to predict light, temperature, and oxygen levels both inside and outside coral tissue; however, this model neglected the influence of ciliary beating and also adopted a constant photosynthetic/respiratory rate [35].

Although the above discussed 2-D or 3-D models are capable of predicting the spatial distribution of oxygen as well as other environmental variables, they all assumed either a constant oxygen generation flux or a constant oxygen concentration on coral surface. In reality, this rate is determined by rates of both photosynthesis and respiration taken place inside the coral. These two important reactions are influenced by the dynamically changing environment, such as temperature and light, as well as carbonate concentration and alkalinity. Moreover, the reactions together with the coupled calcification process can lead to the change of the environment (e.g., pH condition). Consequently, a physically-sound model must take into account all key physiological and carbonate equilibrium reactions, as well as the complex interactions between corals and their environment. Nakamura et al. integrated 3-D ocean hydrodynamic model with the coral polyp physiology model to evaluate how coral calcification at the coral reef scale responds to ocean acidification and sea level rise [36]. Disa et al. further considered the biogeochemical elemental cycling model (BEC) to investigate the interplay between the circulation, seawater carbonate chemistry and the activity of the corals [37]. However, these macroscopic reef-scale framework is not capable of revealing what happens locally at individual polyp scale.

To explore the intricate relationship between dynamically changing external environment and coral health, this study developed a coral-environment system model from a chemical reactor engineering perspective. This model considers photosynthesis, respiration, and calcification reactions within coral tissues, bidirectionally coupled with mass exchange between coral and the surrounding seawater. The exchange of both DO and DIC throughout 24 hours in a day is taken into account. Moreover, ciliary beating that influences localized flow field and hence the mass transport has been integrated into the model. Such a comprehensive model framework enables real-time prediction of coral responses to the changing environmental conditions.

By resorting to this model, we first explored the regulatory role of ciliary beating for coral health. Previous studies have confirmed that ciliary beating on coral surfaces regulates local oxygen concentrations [32-34]. In this study, we delved deeper into examining how ciliary beating controls the exchange of both DO and DIC across the coral-seawater interface, and hence the physiological activities within the coral tissues. Subsequently, we focused on identifying dominant environmental triggers of coral bleaching and underlying mechanisms, particularly global warming and elevated atmospheric carbon dioxide levels were systematically investigated. We discussed their respective impacts on corals during daytime and nighttime in order to identify the underlying causes of symbiotic relationship disruption. It is anticipated that these interesting findings will offer crucial mechanistic insights into coral bleaching and promote the development of effective conservation strategies for reef ecosystems.

## 2. Methodology

### 2.1 Coral geometry in the model

Corals can be generally categorized into two primary groups: hard corals and soft corals [38], or alternatively, warm-water corals and cold-water corals [39, 40]. The former classification is based on the presence or absence of a massive rock-like skeleton, while the latter depends on whether the corals engage in symbiosis with zooxanthellae [41]. This study specifically focuses on hard corals under warm-water environment, as these organisms are the predominant victims of large-scale coral bleaching events.

Hard corals exhibit a variety of morphological forms, among which staghorn coral stands out as an important representative, characterized by its branching structure similar to a stag*’*s antlers (see Fig. 2a). Our modeling domain is centered on the cross-section of a single branch of staghorn coral, along with its adjacent seawater (Fig. 2b). This selection enables us to comprehensively explore the intricate physiological activities taken place within the coral tissue, as well as its dynamic interaction with the surrounding environment. Note that the overall environment is illustrated in Fig. 1a, where the air is also included into the modeling system. As illustrated in Fig. 2c, the cross-section of coral is idealized as a perfect circle, which is composed of coral tissue and skeleton arranged from the outer to the inner regions.

**Figure 2.**
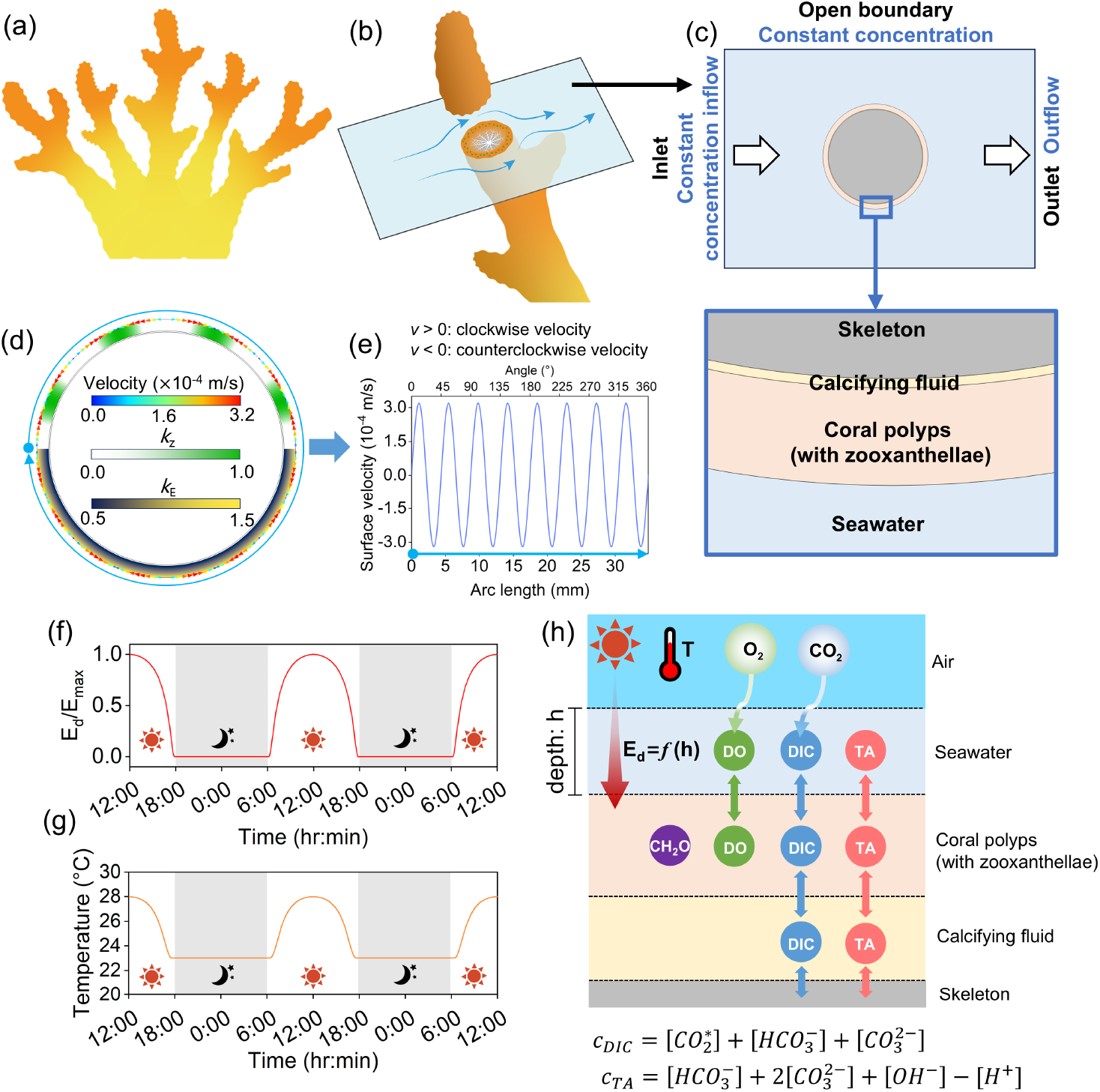
Coral geometry diagram, as well as the computational domain and parameter settings in the model. (a) Morphology of a staghorn coral. (b) Computational domain configuration: rectangular box with seawater flowing around a coral*’*s cross-section. (c) Boundary conditions and layered domain structure: seawater, coral polyp tissue, calcifying fluid, and skeleton (respectively from outer to inner layers). Enlarged view highlights interfacial details. (d) Boundary velocity profile induced by ciliary beating (outer surface) and internal parameter distributions: *k*_*E*_ in Equation (24) and *k*_*z*_ in Equation (25). (e) Coral surface velocity magnitude derived from Equation (1). (f and g) Diurnal-nocturnal evolutions of environmental parameters: light intensity (Equation (14)) and temperature (Equation (15)) profiles over a 2-day cycle. (h) Illustration of domain-specific materials and cross-domain mass transfer. Temperature homogeneity is assumed across all domains, while light intensity exhibits distance-dependent attenuation.

The coral tissue is the ring-shaped envelope that covers the biomineralized skeleton [25, 42]. Coral tissue is composed of two principal layers. The layer adjacent to the seawater is designated as the coral polyp layer, which accommodates symbiotic zooxanthellae capable of conducting both photosynthesis and respiration. The second layer, in contact with the skeleton, is known as the calcifying fluid layer; it establishes a conducive environment for calcification reactions on the skeleton*’*s surface. The skeleton is assumed to be an ideal circle with a diameter of 10 mm. Both layers are assumed to be ideally circular rings with uniform thickness: *d*_*pol*_ = 500 μm for the coral polyp layer and *d*_*cal*_ = 50 μm for the calcifying fluid layer. Additionally, the external seawater environment is modeled as a rectangle to enable a directional seawater flow field.

### 2.2. External environmental condition

#### 2.2.1. Seawater flow and Ciliary beating

The seawater flow surrounding the coral is assumed to be unidirectional, with seawater entering from the left side of the rectangular domain and exiting from the right side (Fig. 2c). The upper and lower boundaries of the rectangle are treated as open boundaries. Given the seawater flow velocity and coral size specified in this study, the estimated Reynolds number is around 2 to 3, indicating laminar flow conditions.

The cilia on coral*’*s surface are capable of beating, thereby disturbing the microenvironment near the coral surface (see Fig. 1a, b). In this study, according to the validated beating models proposed by Shapiro et al. and Pacherres et al., ciliary beating is modeled as the surface velocity *v*_*s*_(m s^−1^) at the coral-seawater interface [32, 34], which follows the next equation:

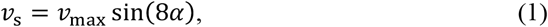

where *v*_*max*_ is the maximum surface velocity (m/s); α denotes the angle (°), which spans from 0 to 360 degrees, as indicated by the blue arrow encircling the coral in Fig. 2d. The calculated surface velocity is depicted by the triangular arrows, where the color corresponds to the velocity magnitude and the orientation indicates the velocity direction. The direction of boundary velocity alternates cyclically between clockwise and counter-clockwise along the coral surface. The specific magnitude of the velocity along the arc is illustrated by the curve in the Fig. 2e, where positive and negative values indicate clockwise and counter-clockwise velocity directions, respectively.

The velocity field in the seawater region is determined by solving the following momentum transfer and continuity equations:

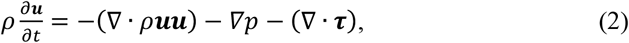

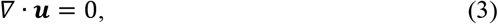

where *u* is the velocity vector (m s^−1^); *t* is the time (s); *p* is the pressure (Pa); *τ* is the stress tensor (N m^−2^); *ρ* is the seawater density (kg m^−3^), assuming uniformity.

#### 2.2.2. Carbonate equilibrium

Atmospheric carbon dioxide dissolved in water transforms into various carbonates. The carbonate equilibrium in both seawater and coral holds significant importance for coral*’*s physiological processes. The series of equilibria representing the reaction when carbon dioxide dissolves in water is as follows:

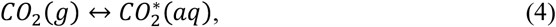

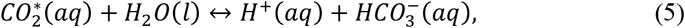

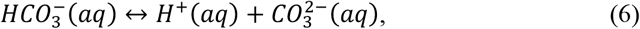

where the symbols (*g*), (*l*), and (*aq*) denote the physical states of the species, corresponding to gaseous, liquid, and dissolved solute states, respectively. 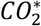 is the state where atmospheric carbon dioxide dissolves in seawater, including both carbon dioxide and carbonic acid. The equilibrium relationships among the concentrations of these various species can be expressed as:

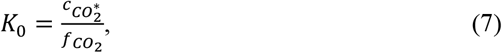

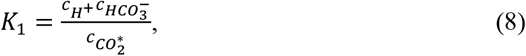

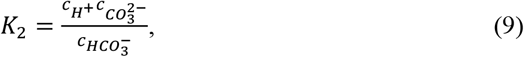

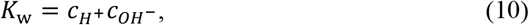

where *c*_*k*_ represents the concentration of each substance indicated by its subscript 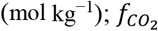 is the fugacity of carbon dioxide in the atmosphere (Pa), which is a function of the volume fraction of atmospheric carbon dioxide 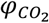, atmospheric pressure, and temperature; *K*_0_, *K*_1_, *K*_2_ and *K*_w_ are equilibrium constants that vary with the temperature and salinity of seawater [43]. Therefore, 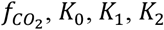, *K*_0_, *K*_1_, *K*_2_ and *K*_w_ will all change simultaneously in response to temperature variations.

The concentrations of the five chemical substances in Equations (7-10) (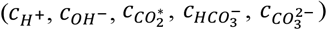) can be organized into two parameters, namely total dissolved inorganic carbon (DIC) and total alkalinity (TA):

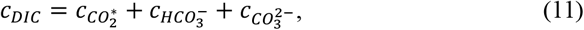

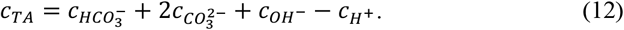

The above six Equations (7-12) are used to describe the carbon dioxide system in the ocean. In our model, they are first employed to determine boundary conditions (*c*_*DIC*_ and *c*_*TA*_), which essentially define the characteristics of the bulk marine environment. This model assumes that the *c*_*TA*_ remains constant in the seawater environment. Consequently, a constant concentration was set at the boundaries, encompassing both the inlet and the open boundaries on either side. The *c*_*DIC*_ at these boundaries can be calculated according to Equations (7-12), which will vary under day-to-night temperature changes.

The *c*_*TA*_ and *c*_*DIC*_ in the seawater domain are calculated using mass-transfer equation based on the boundary concentration conditions mentioned above, in conjunction with the fluxes of TA and DIC across the coral-seawater interface generated by coral physiological activities. Detailed description of physiological reactions within corals can be seen in section 2.3.

#### 2.2.3. Light intensity and Seawater temperature

Both light intensity and seawater temperature determine the efficiency of photosynthesis and respiration of zooxanthellae within coral tissues. The simulation time frame of this study spans 2 days, necessitating the consideration of variations in light intensity and temperature during the day-night transition. The intensity of light reaching the coral surface undergoes attenuation in both the atmosphere and seawater, with the transmission path altering in accordance with changes in solar altitude. The diurnal variation of light intensity can be calculated using Beer*’*s law:

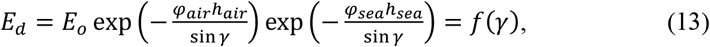

where *E*_*d*_ is the photon flux density reaching the coral surface (mol·m^−2^·s^−1^); *E*_*o*_ is the initial photon flux density emitted from the sun (mol·m^−2^·s^−1^); *h*_*air*_ and *h*_*sea*_ are the thickness of the atmosphere and the depth of the ocean where the coral is located (m), respectively; *φ*_*air*_ and *φ*_*sea*_ are the attenuation coefficients of sunlight by the atmosphere and ocean (m^−1^), respectively, assuming spatial uniformity; *γ* is the solar altitude (°), which is the sole variable in Equation (13) that changes during the day and remains zero at night. The diurnal variation of the solar altitude depends on the local latitude and date. We assume that the coral in the model is located at the equator during an equinox. Consequently, the solar altitude will vary linearly from 0° to 90° and back to 0° throughout the day. The maximum irradiance reaching the coral occurs when the solar altitude angle is 90°, corresponds to 12:00 o*’*clock noon time:

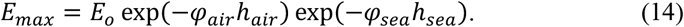

The diurnal variation curve of the final normalized solar irradiance 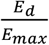 reaching the coral surface is depicted in the Fig. 2f. It increases from 6:00 to 12:00 and decreases from 12:00 to 18:00, while solar radiation is considered to be absent during 18:00 to the next morning 6:00.

Regarding temperature, this study assumes that the diurnal variation trend of temperature *T* (℃) aligns with that of illumination, and the specific formula is:

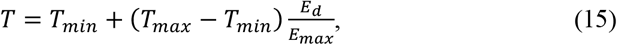

where *T*_*min*_ and *T*_*max*_ are the lowest and highest temperatures of the day (℃). The specific waveform curve is shown in Fig. 2g.

### 2.3. Reactions inside the coral reactor

Coral and zooxanthellae engage in various physiological activities to maintain survival and growth. The photosynthesis of algae, calcification of corals, and respiration of both are the three most critical reactions:

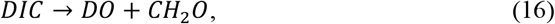

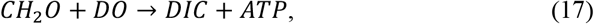

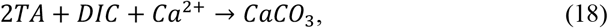

where *DO, CH*_2_*O* and *ATP* represent dissolved oxygen, stored organic matter and adenosine triphosphate respectively. Reaction rates are controlled by the following equations [28]:

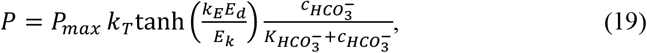

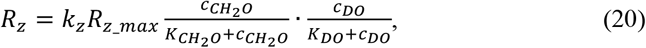

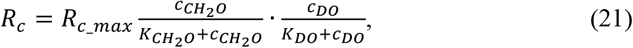

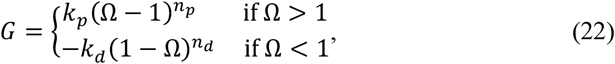

where *P* is the photosynthetic rate of symbiotic algae (mol m^−3^ s^−1^), controlled by light irradiance *E*_*d*_, temperature *T* and bicarbonate ions concentration 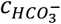, which can be calculated through carbonate equilibrium; *k*_*T*_ is a parameter representing the influence of temperature on photosynthesis, and the formula is as follows [44]:

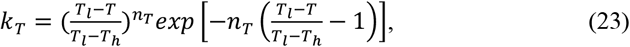

where *T*_*l*_ denotes the upper temperature limit where photosynthetic rate is lowest, while *T*_*h*_ represents the optimal temperature for highest rate (°C); *n*_*T*_ is a dimensionless parameter for sensitivity.

The intensity of light reaching the surface of coral undergoes attenuation in the atmosphere and seawater, and exhibits diurnal variations. Moreover, the distribution of light irradiance within coral tissue is non-uniform. The surface area experiences higher irradiance due to multiple scattering by the skeleton [45], which gradually decreases with increasing tissue depth [27]. The parameter *k*_*E*_ in Equation (19) is employed to account for the depth-dependent variation of light intensity inside the coral:

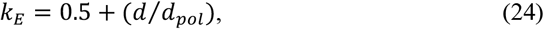

where *d* is the distance from the coral polyps-calcifying fluid interface, and *d*_*pol*_ is the total depth of the polyp layer (m). This equation implies that the surface of the coral in contact with seawater is exposed to light intensity 1.5 times that of the reference, which gradually decreases to 0.5 times the reference intensity as it reaches the calcifying fluid interface.

*R*_*z*_ and *R*_*c*_ in Equations (20-21) are the respiratory rates of zooxanthellae and coral polyps (mol m^−3^ s^−1^), respectively. These rates are regulated by the concentrations of DO and stored organic matter (*c*_DO_and 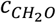). Additionally, *R*_*z*_ is also dependent on the concentration of zooxanthellae and is adjusted by the parameter *k*_*Z*_:

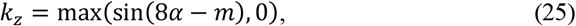

where *α* is the angle (°), consistent with Equation (1); *m* is a parameter used to adjust the distribution of algae within coral polyps (°). Here, *m* is taken as 90° to match experimental observations [34]. The specific distributions of the parameters *k*_*z*_ and *k*_*E*_ are illustrated in the upper and lower regions of the coral polyps, respectively, as depicted in Fig. 2d.

*G* in Equation (22) is the calcification rate (mol m^−2^ s^−1^), which is determined by the aragonite saturation state Ω:

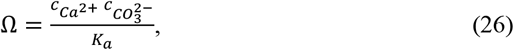

where *K*_*a*_ is the solubility product for aragonite that changes with temperature (mol^2^·kg^−6^) [46]. Given that the variation in 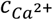 is substantially smaller than that in 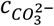, a constant value of 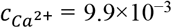 mol kg^−1^ was adopted for the calcifying fluid [28].

Note that the calculation of photosynthetic rate *P* and calcification rate *G* requires the determination of 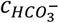 and 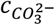, which can be calculated based on the equations in section 2.2.2. Equations (8-12) are employed to compute the values of 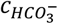 and 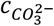 according to the *c*_*TA*_ and *c*_*DIC*_ within the domain.

### 2.4. Mass transfer

The section 2.2.2 has outlined all the species accounted for in our model, however, not all of them are required to be incorporated into mass transfer calculations. The substances considered in each region are illustrated in Fig. 2h (Seawater: DO, DIC, TA; Polyps: DO, DIC, TA, CH_2_O; Calcifying fluid: DIC, TA). Species transport within the computational domain is governed by the following equations:

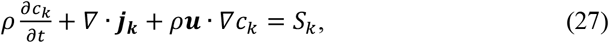

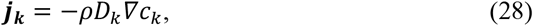

where *c*_*k*_ is the concentration of each substance (mol kg^−1^); ***j***_***k***_ is the diffusive flux (mol m^−2^ s^−1^); *S*_*k*_ is the source term (mol m^−3^ s^−1^), arising from photosynthesis (see Equation (19)), respiration (see Equation (20-21)), and calcification (see Equation (22)); *D*_*k*_ is the mass diffusion coefficient of each substance (m^2^ s^−1^). A fixed value of mass diffusion coefficient is assigned to DIC and TA related species, i.e., the diffusion coefficient of carbonate ion [47]. The stored organic matter is assumed as glucose, so the mass diffusion coefficient of CH_2_O is set as the diffusion coefficient of glucose [48].

The mass exchange across seawater-coral interface is driven by concentration gradient and also controlled by the above equations. But the exchange between coral polyps and calcifying fluid is regulated by the energy generated through respiration, as well as the paracellular pathway. The former is calculated using the following formula [28]:

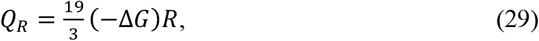

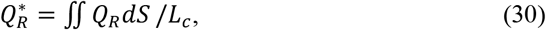

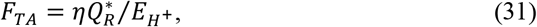

where *Q*_*R*_ is the energy produced by respiration (J m^−3^ s^−1^), specifically from the hydrolysis of ATP; Δ*G* is the free energy released by ATP hydrolysis (J mol^−1^); Owing to the heterogeneous spatial distribution of zooxanthellae, respiratory activity exhibits significant regional variation within coral tissues. This study assumes that the energy generated by respiration is evenly applied to the polyps-calcifying fluid interface, ignoring the issue of inhomogeneous energy generation. Consequently, an average energy 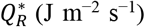 on the interface was calculated by integrating the *Q*_*R*_ over the coral polyp and dividing it by the interface length *L*_*c*_ (i.e., Equation (30)).

*F*_*TA*_ is the TA flux from coral polyps to the calcifying fluid (mol m^−2^ s^−1^); *η* is the energy conversation efficiency; 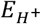 is the energy required to transport 1 mol of *H*^+^ (J/mol), which is equivalent to the energy needed to transport 1 mol of TA. The value of 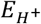 depends on the pH difference and the 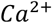 concentration difference between coral polyps and the calcifying fluid:

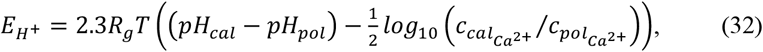

where *R*_*g*_ is the gas constant (J mol^−1^ K^−1^); *T* is the Kelvin temperature (K). The 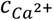 difference is significantly smaller than the pH difference, and the daily variation in 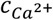 is also much less pronounced compared to that of pH. Therefore, the term 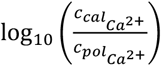 can be approximated as a constant (∼0.02) [28]. Consequently, Equation (32) is rewritten as follows:

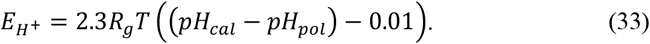

Beyond energy-driven active transport (i.e., Equations (29-33)), TA can also undergo passive transport through paracellular pathway, and these pathways similarly permit DIC passage [28]:

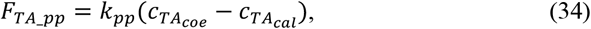

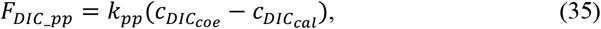

where *F*_*TA_pp*_ and *F*_*DIC_pp*_ are the flux from coral polyps to the calcifying fluid through paracellular pathway (mol m^−2^ s^−1^); *k*_*pp*_ is conductivity coefficient for the paracellular leakage process (m/s).

## 3. Results and discussion

### 3.1. Exploring the functions of coral cilia beating

The ciliary array covering the coral surface generates coordinated beating motions, creating special localized flow conditions. Experimental observations indicate that the collective activity of these cilia produces vortices that regulate the DO environment on the coral surface, making it suitable for coral survival [32-34]. By resorting to the chemical reactor engineering model developed in this study, we will further elucidate the function of beating on coral health. All parameter values are listed in the Appendix.

#### 3.1.1. Formation of near-surface vortex field

Our model simulates ciliary beating by implementing boundary velocities along the coral surface. Compared to the case without beating, the dynamic motion of cilia generates multiple near-surface vortices (see the velocity field and streamline distributions in Fig. 3a). Due to these vortices, the flow direction on the coral surface regularly shifts from flowing towards the coral to flowing away from the coral (see the velocity arrows in Fig. 3b, where color indicates the direction of flow, either towards the coral or towards the bulk water). In the absence of ciliary beating, the flow field exhibits typical characteristics of a low-Reynolds-number cylindrical flow, where seawater flows smoothly and steadily over the coral surface without noticeable vortex formation (see the subplots on the right side in Fig. 3a, b).

**Figure 3.**
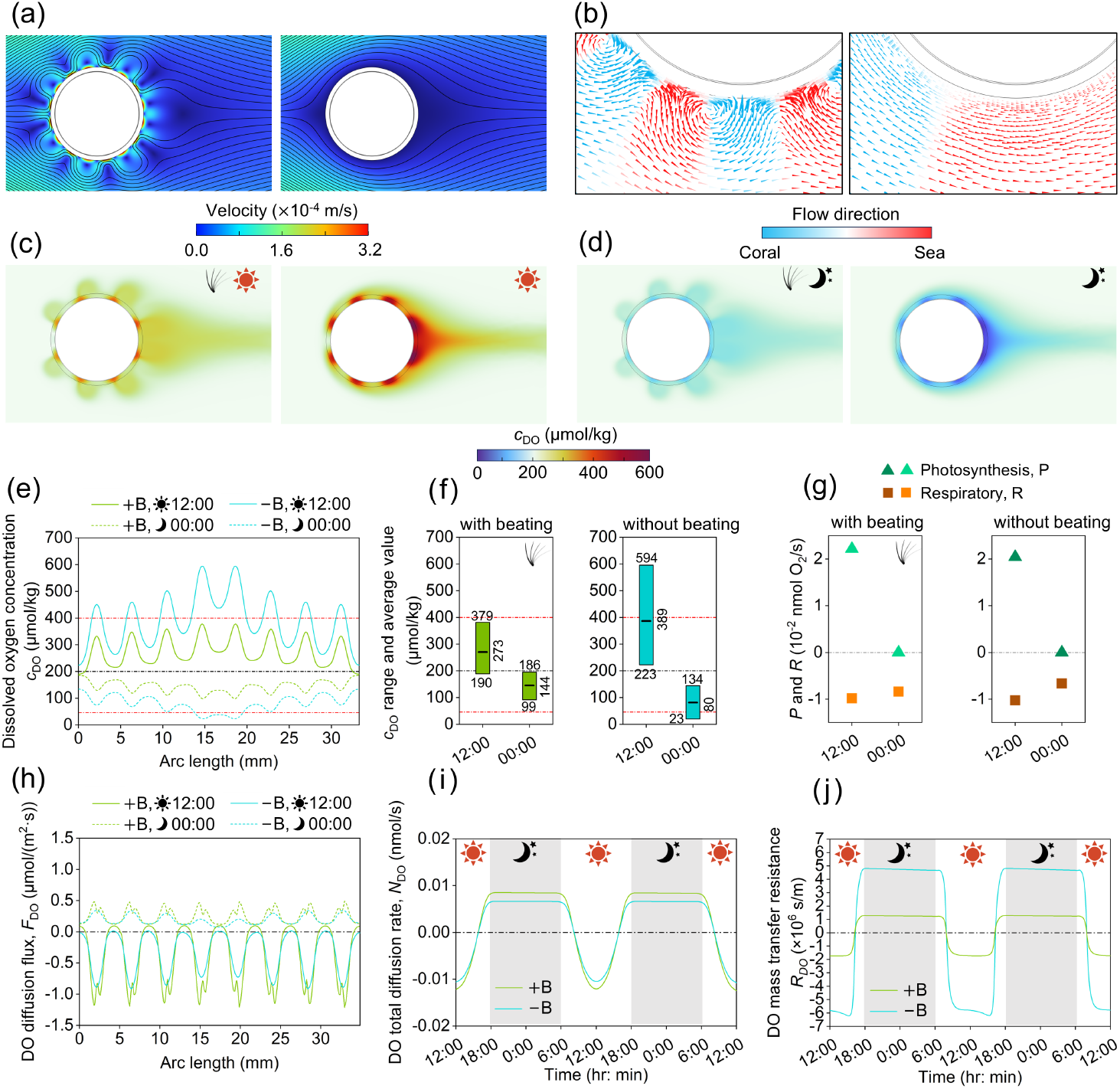
How ciliary beating can regulate dissolved oxygen transport and exchange. (a) Flow field comparison under ciliary beating (left) versus without beating (right), visualized by streamlines (black line) and velocity magnitude (surface color). The color bar is located at the bottom. (b) Local velocity vector distributions: ciliary beating (left) induces vortices formation and cyclic flow directional change (flow to coral: blue; to distal ocean: red), no vortex can be identified for the case without beating (right). The color bar is located at the bottom. (c, d) Spatial distributions of DO concentrations at noon (12:00) (c) and at midnight (00:00) (d) for both cases with and without beating. (c) and (d) share the same color bar located at the bottom. DO concentration evolutions over a two-day time range are given in the Supplementary Animation 1. (e) DO concentration distribution within coral tissue (+B: with beating; −B: without beating). (f) Quantitative analysis of DO concentration range and mean values in coral tissue. (g) Average rate of photosynthesis and respiration in coral tissue. (h) Spatial distribution of DO flux along the coral-seawater interface (positive: inhale, from seawater to coral; negative: exhale, from coral to seawater). (i) Diurnal variation of total DO transfer rate across the coral-seawater interface over two days. (j) Diurnal variations in mass transfer resistance for DO transfer from the coral surface to the distal ocean.

The hydrodynamic changes induced by ciliary activity in the near-surface region significantly affect certain environmental conditions, particularly the availability of oxygen and carbon dioxide. We will address these impacts separately in the following two sections.

#### 3.1.2. Regulation of dissolved oxygen exchange

A fundamental physiological prerequisite for sustaining coral viability lies in the maintenance of appropriate DO concentrations. Both hyperoxic and hypoxic conditions can detrimentally impact coral health: excessive DO leads to the generation of toxic free radicals [9, 16], while DO deficiency directly impairs the respiration of coral polyps and their symbiotic zooxanthellae.

Our comparative analysis demonstrates that the near-surface vortices induced by ciliary activity significantly improves coral living conditions compared to the case without ciliary beating. It is interesting to discover that the vortices generated by cilia regulate DO exchange in a smart way and ciliary beating serves distinctively different functions during the day versus at night. During daylight hours, these vortices prevent the accumulation of high oxygen levels on the coral surface by dispelling oxygen-saturated seawater away from coral surface while drawing in normally oxygenated water (see the DO distribution during the day in Fig. 3c). On the other hand, at night, nocturnal ciliary activity helps alleviate extreme hypoxic conditions. The vortical flow displaces oxygen-depleted seawater surrounding the coral and introduces oxygen-rich seawater from the bulk (see the DO distribution at night in Fig. 3d).

This near-surface regulation enables coral surface seawater to maintain an appropriate DO concentration, which affects the DO concentration within coral tissue. The DO concentrations within coral polyps are recorded in Fig. 3e, where the black dotted line represents the ambient oceanic oxygen level and the red dotted lines indicate the thresholds for hyperoxia and hypoxia. The data collection position is located at a distance of 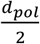 from the seawater-polyp interface; the starting and ending positions are indicated in Fig. 2d. The hyperoxia threshold is set as twice the ambient oceanic DO concentration, while the hypoxia threshold is 59.5 μmol kg^−1^ [49].

Comparative analysis of DO conditions within coral tissues reveals that the cessation of ciliary beating results in a marked elevation of daytime oxygen accumulation and a more pronounced hypoxic state at night, particularly in regions with abundant zooxanthellae. Additionally, external seawater flow further contributes to the localized daytime buildup of oxygen-saturated water (and nighttime accumulation of oxygen-depleted water) for coral tissues situated downstream of ocean currents, thereby intensifying adverse conditions.

Figure 3f employs a bar chart to quantitatively compare the regulatory of ciliary beating on the DO state within coral tissue. The bar range and the black horizontal line inside each bar denoting the DO concentration ranges and mean values within coral polyps, respectively. When ciliary beating is absent, the average DO concentration significantly increases during the daytime, rising from 273 μmol kg^−1^ to 389 μmol kg^−1^, which represents a 42.5% increase, and decreases notably at night, dropping from 144 μmol kg^−1^ to 80 μmol kg^−1^, a 44.4% reduction.

The alterations in oxygen supply conditions impact the respiratory capacity of corals, as depicted in Fig. 3g. During the day, excessive oxygen levels in the absence of ciliary activity resulted in a slight increase in respiration, with the rate rising from – 0.9824×10^−2^ nmol s^−1^ to –1.0254×10^−2^ nmol s^−1^ (a 4.4% increase; note that positive values signify oxygen production, whereas negative values indicate oxygen consumption). Severe hypoxia at night leads to substantial respiratory distress, with the oxygen consumption rate during respiration decreasing from –0.8412 nmol s^−1^ to – 0.6599 nmol s^−1^ (representing a 21.6% decline). Inhibition of respiration may prompt symbiotic zooxanthellae to abandon coral polyps in search of a more favorable environment, leading to coral bleaching. The above findings suggest that abnormal environmental alterations impacting coral cilia beating can indirectly result in the degradation of the coral*’*s living environment, potentially serving as an underlying cause of coral bleaching.

#### 3.1.3. The impact on dissolved oxygen mass transfer resistance

The regulation of the oxygen environment on coral surfaces and within tissues through ciliary beating, as revealed in Section 3.1.2, is actually accomplished by changing the mass transfer efficiency of DO diffusion from the coral surface to the bulk seawater and vice versa. During the day, DO needs to be dispelled away from the coral surface, while at night, it requires replenishment through the seawater-polyp interface. Coral ciliary beating facilitates this process by decreasing the mass transfer resistance of DO, thereby facilitating DO exchange. Oxygen mass transfer efficiency can be quantified by the flux at the coral-seawater interface. Pacherres et al. indirectly calculated oxygen flux based on the concentration gradient near the coral surface, thus sampling position influences the prediction [33].

Our numerical model allows the calculation of DO flux, denoted as *F*_*DO*_ (as illustrated in Fig. 3h), along the complete polyp-seawater interface. Moreover, it allows the integration of this flux to determine the overall oxygen exchange rate, denoted as *N*_*DO*_ (Fig. 3i, a coral thickness of 1 mm was assumed when calculating interface integration). The spatial distribution of DO flux across the coral surface reveals significant regional disparities. In areas densely populated with zooxanthellae, the elevated rates of photosynthesis and respiration result in more vigorous material exchange with the surrounding seawater, which occurs both during the day and at night.

The diurnal fluctuations in the total exchange rate of DO, *N*_*DO*_, demonstrate that corals predominantly exhibit oxygen exhalation behavior during daylight hours. As sunlight intensity gradually diminishes towards nightfall, the reduced oxygen production through photosynthesis requires corals to absorb external oxygen to support their respiratory processes (as evidenced in Fig. 3i, where *N*_*DO*_ shifts from negative to positive values).

Ciliary beating*’*s contribution to promoting uninterrupted oxygen exchange throughout the diurnal cycle is less pronounced than its role in regulating the oxygen state. Only minor different in *F*_*DO*_ and *N*_*DO*_ can be observed between conditions with and without ciliary beating, with the oxygen flux exhibiting only a slight increase under ciliary beating compared to the case without beating. This phenomenon can be explained by the fact that the quantity of oxygen released (or absorbed) from the external environment must closely match the amount produced (or consumed) through photosynthesis and respiration in corals to satisfy physiological balance, while the disparity in photosynthesis and respiration rates with and without ciliary oscillation remains insignificant (see Fig. 3g).

Mass transfer resistance is a quantifiable indicator for evaluating the transport efficiency of substances, and its calculation formula is:

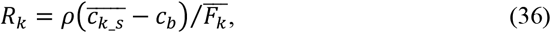

where *R*_*k*_ is the mass transfer resistance of substance 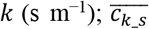 is the average concentration on the coral surface (mol kg^−1^); *c*_*k_b*_ is the concentration in bulk seawater (mol kg^−1^);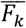 is the average flux on the coral polyps-seawater interface (mol m^−2^ s^−1^).

The quantitative results demonstrate a considerable difference in mass transfer resistance between conditions with and without ciliary beating. Specifically, the mass transfer resistance is significantly higher in the absence of beating, increasing by 3.36 times at noon (Time: 12:00) and 3.76 times at midnight (Time: 00:00) compared to scenarios where beating occurs (see Fig. 3j). This indicates that, under the influence of ciliary beating, oxygen can more easily diffuse from the coral tissue into the surrounding seawater or be obtained from the external environment. In other words, ciliary beating enables corals with stronger mass transfer ability. Although deterioration or lack of this ability may not immediately lead to detrimental sequences, this ability may play a crucial role when corals encounter extreme environmental conditions.

#### 3.1.4. Regulation of dissolved inorganic carbon and pH environment

Existing research efforts on coral*’*s cilia have mainly focused on their role in oxygen regulation, a topic we have elaborated in the preceding section. Their functions beyond oxygen regulation are further explored in this work. Coral cilia also play a crucial role in regulating the concentration of DIC and pH levels within coral tissues.

DIC and pH level are also key environmental parameters for corals. Elevated DIC concentrations typically coincide with reduced pH levels, and the resulting seawater acidification is likely to exert extra stress on coral calcification processes [22]. A lower concentration of DIC, however, means that normal photosynthetic efficiency cannot be maintained, affecting the symbiotic relationship between algae and coral polyps.

The beating of cilia can also effectively regulate the DIC concentration in both coral*’*s surface seawater and tissues, preventing extreme carbon deficiency during the day and excessive accumulation at night (as illustrated in Fig. 4a, b). During daylight hours, ciliary beating facilitates the transport of bulk seawater with elevated DIC concentrations to the coral surface, ensuring an adequate DIC supply within the tissues to sustain photosynthesis. At night, the respirations of both coral polyps and their symbiotic algae release DIC, and ciliary beating promotes efficient dispel of the accumulated DIC away from coral to prevent excessive acidification.

**Figure 4.**
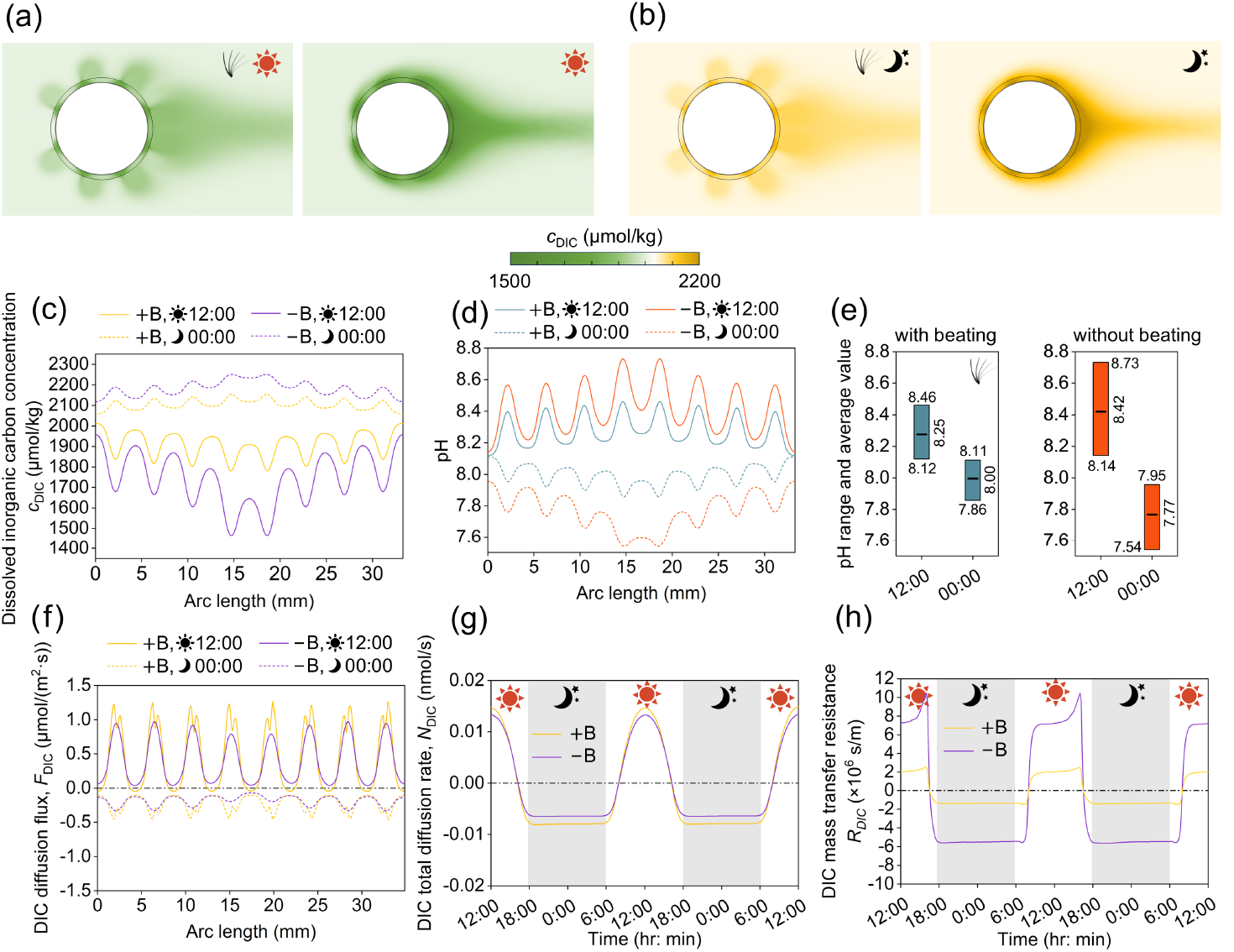
How ciliary beating can regulate pH and the transport and exchange of dissolved DIC. (a, b) Spatial distributions of DIC concentrations at noon (12:00) (a) and at midnight (00:00) (b) for both cases with and without beating. (a) and (b) share the same color bar located at the bottom. DIC concentration evolutions over a two-day time range are given in the Supplementary Animation 2. (c, d) DIC concentration and pH distributions within coral tissue (+B: with beating; −B: without beating). (e) Quantitative analysis of pH values in coral tissue. (f) Spatial distribution of DIC flux along the coral-seawater interface (positive: from seawater to coral; negative: from coral to seawater). (g) Diurnal variation of DIC exchange rate at the coral-seawater interface over 2 days. (h) Diurnal variations in mass transfer resistance for DIC diffusion from the coral surface to the bulk seawater.

The concentration distribution of DIC within coral tissue indicates that the most severely impacted areas are those densely populated with zooxanthellae, where photosynthesis and respiration respectively lead to a higher degree of inorganic carbon consumption and production (as illustrated in Fig. 4c). Since DIC serves as the photosynthetic raw material for zooxanthellae, the absence of beating slightly inhibits photosynthetic efficiency during the day. The daytime photosynthesis rate will be slightly lower, decreased from 2.2175 nmol s^−1^ to 2.0431 nmol s^−1^ (a 7.9% decline, see Fig. 3g).

An abnormal DIC environment can result in unfavorable pH values within coral tissues, with regions of extreme pH also aligning with areas packed with zooxanthellae, where extreme DIC concentrations show up (Fig. 4c,d). The absence of ciliary beating leads to an increase in the average pH value of coral tissue from 8.25 to 8.42 during the day and a decrease from 8.00 to 7.77 at night (Fig. 4d, e).

#### 3.1.5. The impact on mass transfer resistance of dissolved inorganic carbon

The regulatory effect of ciliary beating on DIC can also be quantitatively evaluated by examining the flux *F*_*DIC*_ and exchange rate *N*_*DIC*_ across the polyp-seawater interface, as well as the mass transfer resistance related to its diffusion into the surrounding seawater, similar to the evaluation of DO in section 3.1.3. During daytime, corals predominantly uptake DIC to sustain photosynthetic activity of their symbiotic algae. However, at night, the respirations of coral polyps and their symbiotic zooxanthellae cause the release of carbon dioxide, continuously generating DIC. The effect of ciliary beating remains relatively insignificant on DIC exchange flux at the coral-water interface, merely resulting in a slight augmentation (see the spatial distribution of the DIC flux *F*_*DIC*_ and the diurnal variation of total exchange rate *N*_*DIC*_ at the seawater-polyp interface in Fig. 4f, g, respectively). Nevertheless, from the perspective of mass transfer resistance, it becomes apparent that DIC can be more readily transported between coral surfaces and bulk seawater for the case with ciliary beating. Specifically, ciliary beating significantly decreases the mass transfer resistance of DIC transfer from the bulk seawater to the tissue surface, as evidenced by a 3.56-fold reduction at noon (Time: 12:00). Similarly, the mass transfer resistance governing the back-diffusion of DIC from the tissue surface to the ocean is reduced by 4.01-fold at midnight (Time: 00:00), as depicted in Fig. 4h.

### 3.2. Exploration of the causes of coral bleaching under global warming

A widely reported view on the causes of coral bleaching is the high temperature seawater caused by global warming, combined with high irradiance, can trigger the production of harmful reactive oxygen species (ROS) and reactive nitrogen species (RNS) in coral tissues [9]. Additionally, the concurrent rise in atmospheric carbon dioxide concentration can also impact the physiological activities of corals, including respiration, photosynthesis, and calcification [18]. However, the in-depth mechanism by which these two major global environmental changes induce coral bleaching remains an open question, and it is challenging to uncouple the influences from two factors and to determine whether high temperature or rising atmospheric carbon dioxide is more likely to cause coral bleaching.

To answer these questions, we designed one control and three hypothetic cases. The control case is under normal conditions, where seawater temperature is *T*_*max*_=25 ℃-*T*_*min*_=20 ℃ and atmospheric concentration is 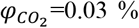 The three abnormal cases are: (1) temperature alone is increase to *T*_*max*_=28 ℃-*T*_*min*_=23 ℃ (at this point, *T*_*max*_ corresponds to the optimal photosynthetic temperature *T*_*h*_); (2) atmospheric carbon dioxide concentration alone is increased to 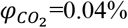 and (3) both temperature and atmospheric carbon dioxide concentration are increased simultaneously. The magnitudes of these changes are based on observations in the South China Sea over the past 50 years [50].

We monitored changes in DO concentration and pH within corals under different scenarios. The DO concentrations in tissues for four different cases are shown in Fig. 5a. Compared to an increase in atmospheric carbon concentration, a temperature rise led to a higher average DO concentration in coral tissue during the day (as illustrated in Fig. 5b). The former increased the average DO concentration by 0.59%, while the latter increased it by 5.18% and caused local areas to exceed the oxygen threshold. The synergistic effect of the two factors increased the average DO concentration by 5.98%, which was slightly higher than the sum of the effects of the first two alone (i.e., 5.98%>5.77%=0.59%+5.18%). Both high-temperature and high atmospheric carbon dioxide concentration had a relatively minor impact on oxygen availability at night.

**Figure 5.**
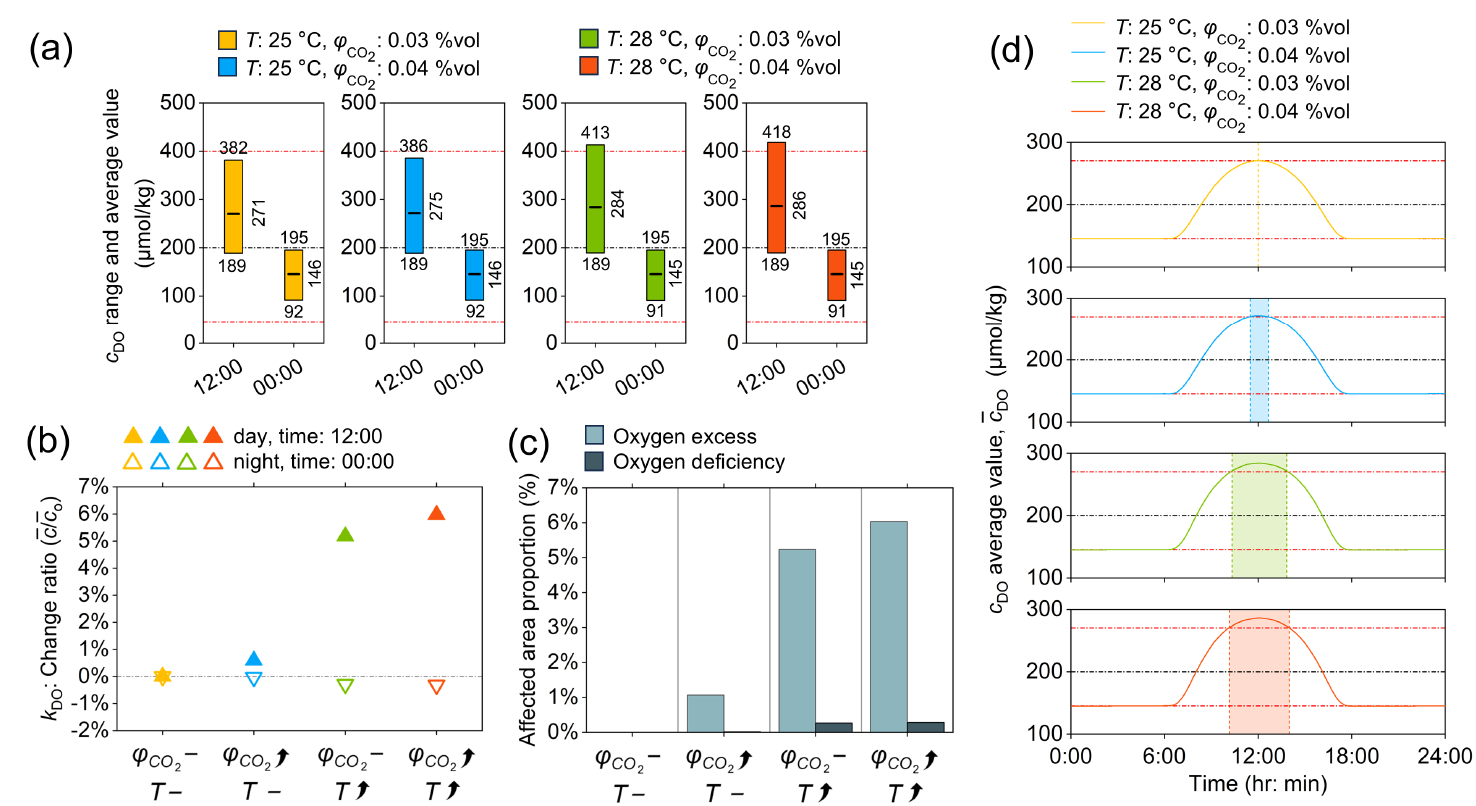
Comparison of coral living environment under different scenarios, focusing on DO. (a) DO concentration range and mean values in coral tissue. (b) Relative increase of the mean value of DO concentration in coral tissue under three stressed conditions compared to the normal condition. (c) Proportion of coral tissue in DO excess/deficiency states under different environmental perturbations. (d) Temporal exposure duration to DO excess/deficiency states for different scenarios.

Figure 5c illustrates the proportion of areas within coral tissue experiencing excess oxygen (defined as DO concentration exceeding the highest value under normal conditions) and hypoxia (defined as DO concentration below the lowest value under normal conditions) in coral tissue for each case. It is evident that the temperature increase has resulted in a larger area of excess oxygen, while also causing small-area hypoxia.

The diurnal variation (0:00-24:00) of the average DO concentration within coral tissue is depicted in Fig. 5d, with the colored regions representing the periods of oxygen excess. The findings reveal that oxygen excess predominantly occurs at noon. Furthermore, in comparison to the increase in DIC concentration, a temperature rise leads to a more prolonged duration of the oxygen excess state.

Compared to the elevated temperature, the increase in atmospheric carbon dioxide concentration has a relatively smaller impact on the DO concentration in corals. But it mainly leads to an increase in DIC concentration in the ocean, which in turn affects the pH value of coral tissue (Fig. 6a). The comparison results in Fig. 6b reveal that the rise in atmospheric carbon dioxide causes the average pH within coral tissues to decrease by 1.11% during the day and 1.66% at night (equivalent to hydrogen ion concentration changes of 25.9% and 38.0%, respectively).

**Figure 6.**
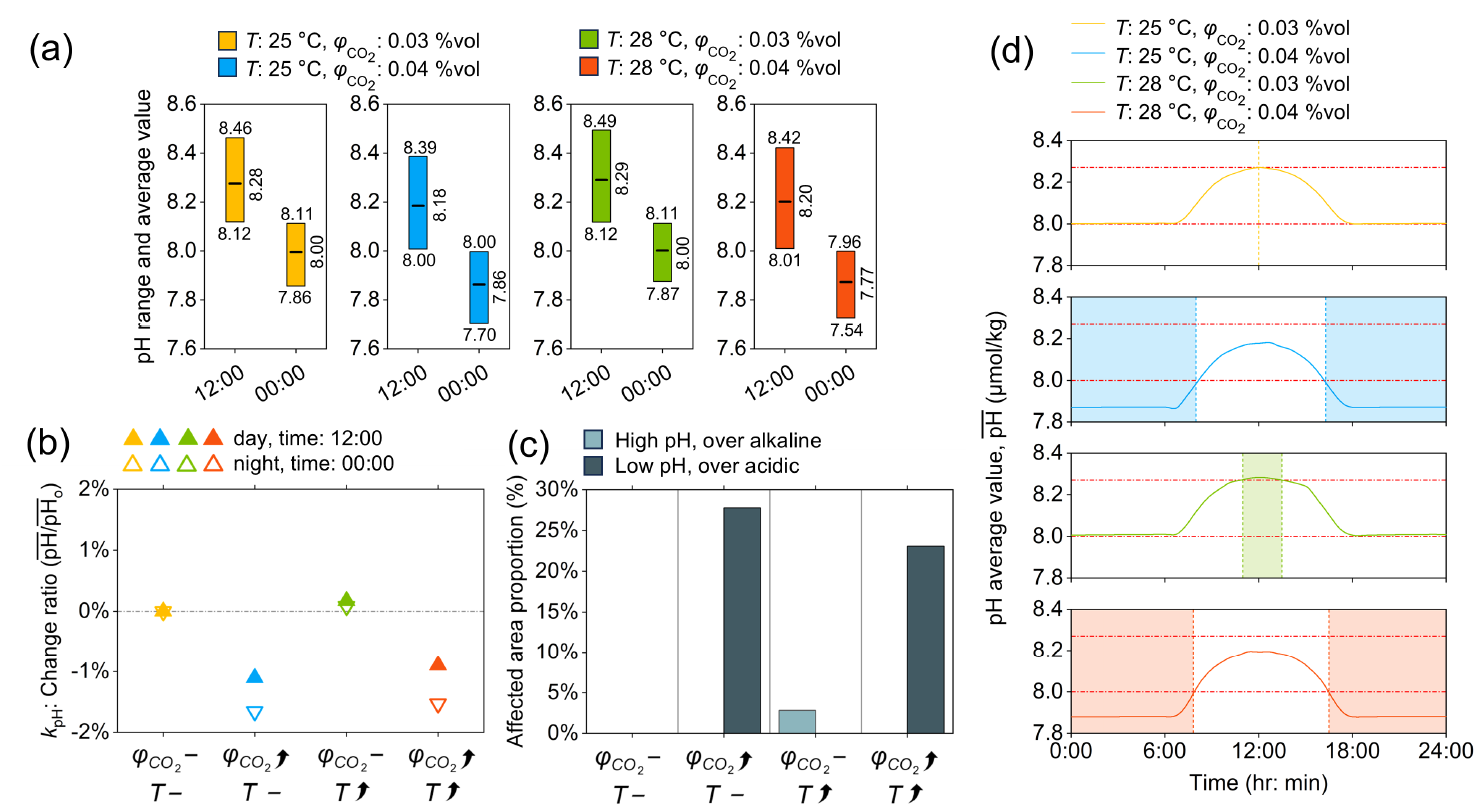
Comparison of coral living environment under different environmental conditions, focusing on pH condition. (a) mean value and range of pH in coral tissue. (b) Relative increase of mean pH in tissue under three stressed conditions compared to the normal condition. (c) Proportion of coral tissue in high/low pH states under different environmental perturbations. (d) Temporal exposure duration to high/low pH states for different scenarios.

During the day, coral experiences a higher pH due to the consumption of DIC through photosynthesis. Therefore, the daytime pH decrease does not result in areas being affected by acidity. At night, however, due to carbon production from respiration, the pH is lower, and the increase in atmospheric carbon dioxide further intensifies the acidic environment. The proportion of areas with acidity (defined as pH lower than the lowest value under normal condition) at night reaches as high as 27.8%, and this condition persists throughout the whole night (see Fig. 6d).

The temperature increase has minimal impact on the pH value within coral tissues, causing only a slight increase in the average pH value (1.71% during the day and 0.81% at night, as illustrated in Fig. 6b), with a smaller fraction of areas and a shorter duration affected by alkalinity (as illustrated in Fig. 6c, d). Therefore, overall, the increase in atmospheric carbon dioxide concentration primarily affects corals by creating an acidic tissue environment at night, leading to algal intolerance.

Comparative analyses of the simulation results of four cases tell us the primary impacts of increased seawater temperature and atmospheric carbon dioxide levels on coral health, driven by global warming. Elevated temperature predominantly exerts stress during daylight hours, augmenting DO concentrations through enhanced photosynthesis, which in turn accelerates the production of detrimental reactive oxygen species (ROS). Concurrently, increased atmospheric carbon dioxide levels mainly contribute to seawater acidification, with a pronounced effect observed particularly at night. During night, the respiration and DIC production by corals and their symbiotic algae intensify the acidification process, further deteriorating the acidification state of the coral habitat.

## 4. Summary and outlook

This study develops a comprehensive coral-environment system model from a chemical reactor engineering perspective. This model takes into account multiple reaction units, i.e., coral polyps and their algal symbionts, where key physiological processes such as photosynthesis, respiration, and calcification take place. The model also incorporates bidirectional mass exchange between the bioreactor (i.e., coral) and its external environment (i.e., seawater), and among coral*’*s internal reaction units. The exchanged substances include dissolved oxygen, carbon species, and other metabolically essential compounds.

Another unique feature of the model is the inclusion of ciliary beating, which generates localized near-surface vortices to enhance mass transfer efficiency. This flow pattern was rigorously analyzed in this study to quantify its role in dispelling locally accumulated *“*harmful*”* species (e.g., supersaturated DO during daytime and DIC at night) by reducing diffusion resistance. Such mechanisms can offer favorable intracellular conditions for physiological activity and stress resilience.

Thanks to the consideration of the interaction between the coral and its external environment, global warming-induced seawater temperature increase and atmospheric carbon dioxide elevation could be separately evaluated to uncouple their distinct impacts. Elevated temperature primarily enhances daytime photosynthetic efficiency, increasing DO concentration in coral tissue, which leads to the risk of overproduction of reactive oxygen species (ROS) and reactive nitrogen species (RNS). Elevated carbon dioxide, on the other hand, acidifies nocturnal tissue microenvironments, potentially disrupting symbiont tolerance and calcification kinetics.

Compared to existing models, this work stands out in modeling coral–environment interactions by implementing coral-specific physiological features (including physiological reactions and ciliary-driven flow) and considering diurnal environmental variability (light, temperature, and pH). This general model, refined through gradual integration of biological and physical features for specific coral systems, will be a powerful predictive tool that can guide conservation strategies, and help coral ecosystems tackle the escalating impacts of climate change.

## Supporting information

Supplementary Animation 1

Supplementary Animation 2

## Acknowledgements

We are grateful for the financial support from the National Natural Science Foundation of China (21978184), the *“*Jiangsu Innovation and Entrepreneurship (Shuang Chuang) Program*”*, the *“*Jiangsu Specially Appointed Professors Program*”*, and the *“*Priority Academic Program Development (PAPD) of Jiangsu Higher Education Institutions*”*. Mr. Jian Han*’*s preliminary efforts on this project are also appreciated.

## Appendix

**Table A1.**
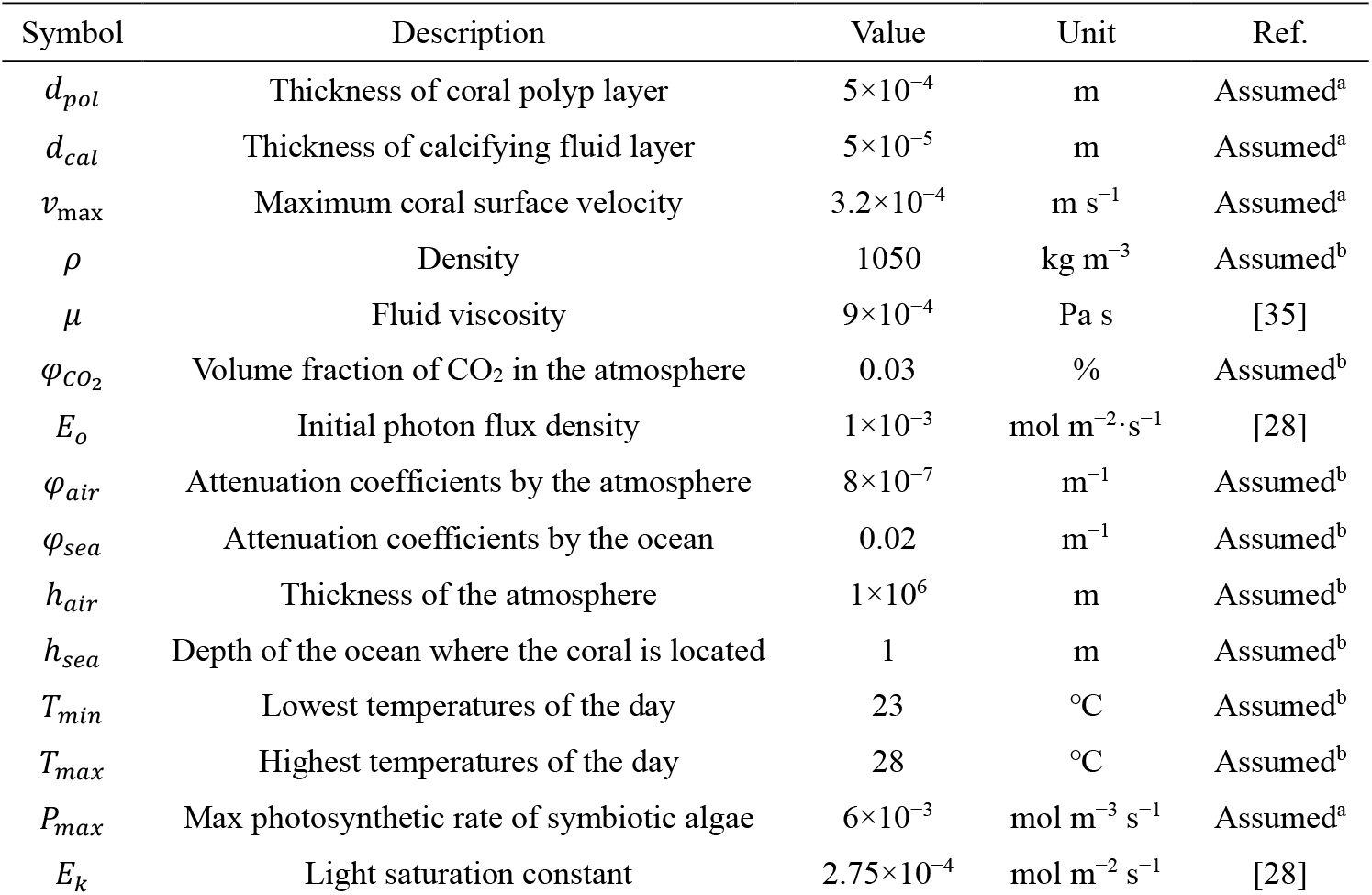
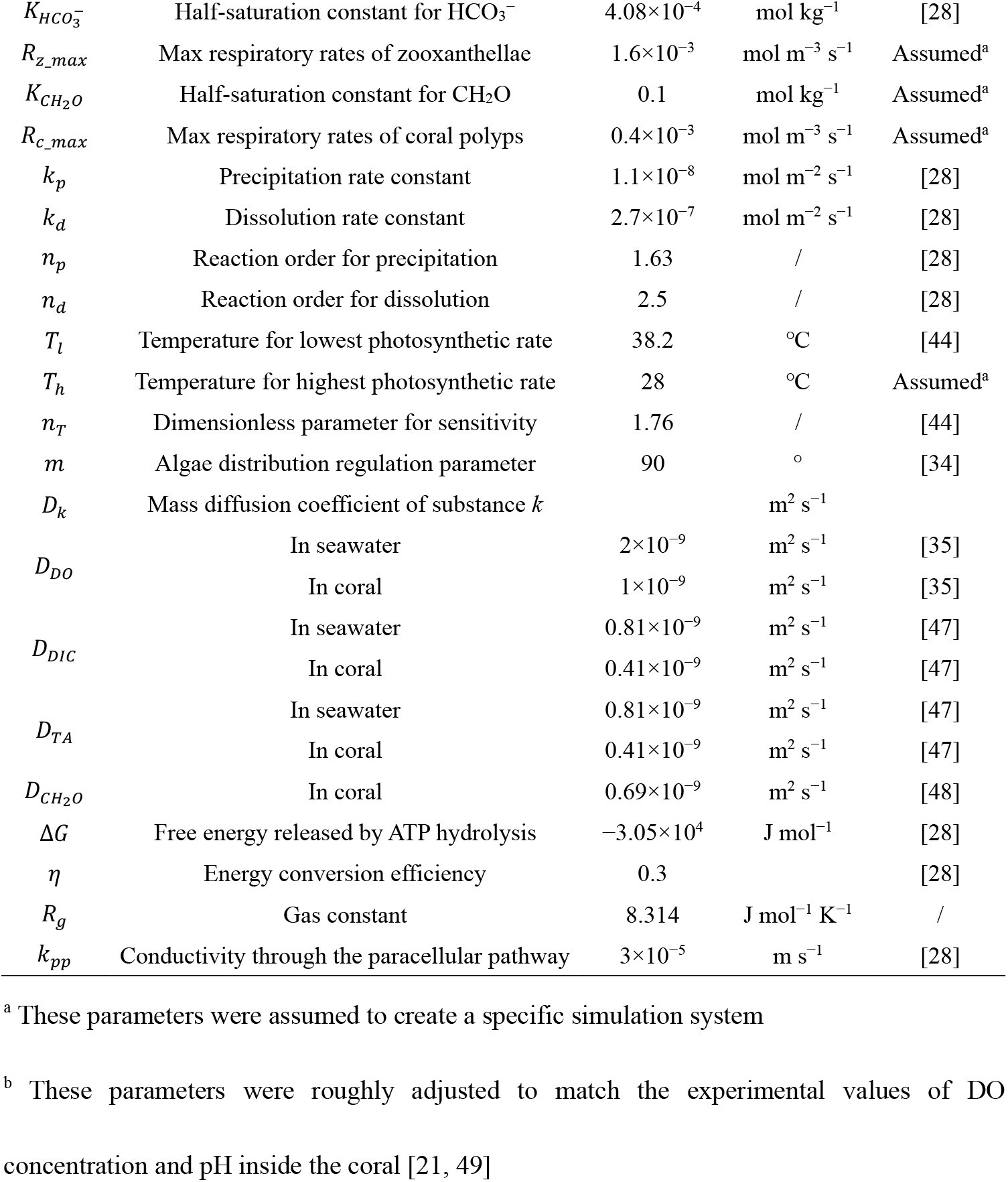
The parameters with constant values (for the control case)

| Symbol | Description | Value | Unit | Ref. |
| --- | --- | --- | --- | --- |
| $d_{pol}$ | Thickness of coral polyp layer | $5 \times 10^{-4}$ | m | Assumed<sup>a</sup> |
| $d_{cal}$ | Thickness of calcifying fluid layer | $5 \times 10^{-5}$ | m | Assumed<sup>a</sup> |
| $v_{max}$ | Maximum coral surface velocity | $3.2 \times 10^{-4}$ | $\text{m s}^{-1}$ | Assumed<sup>a</sup> |
| $\rho$ | Density | 1050 | $\text{kg m}^{-3}$ | Assumed<sup>b</sup> |
| $\mu$ | Fluid viscosity | $9 \times 10^{-4}$ | $\text{Pa s}$ | [35] |
| $\varphi_{CO_2}$ | Volume fraction of CO<sub>2</sub> in the atmosphere | 0.03 | % | Assumed<sup>b</sup> |
| $E_o$ | Initial photon flux density | $1 \times 10^{-3}$ | $\text{mol m}^{-2} \cdot \text{s}^{-1}$ | [28] |
| $\varphi_{air}$ | Attenuation coefficients by the atmosphere | $8 \times 10^{-7}$ | $\text{m}^{-1}$ | Assumed<sup>b</sup> |
| $\varphi_{sea}$ | Attenuation coefficients by the ocean | 0.02 | $\text{m}^{-1}$ | Assumed<sup>b</sup> |
| $h_{air}$ | Thickness of the atmosphere | $1 \times 10^6$ | m | Assumed<sup>b</sup> |
| $h_{sea}$ | Depth of the ocean where the coral is located | 1 | m | Assumed<sup>b</sup> |
| $T_{min}$ | Lowest temperatures of the day | 23 | °C | Assumed<sup>b</sup> |
| $T_{max}$ | Highest temperatures of the day | 28 | °C | Assumed<sup>b</sup> |
| $P_{max}$ | Max photosynthetic rate of symbiotic algae | $6 \times 10^{-3}$ | $\text{mol m}^{-3} \text{s}^{-1}$ | Assumed<sup>a</sup> |
| $E_k$ | Light saturation constant | $2.75 \times 10^{-4}$ | $\text{mol m}^{-2} \text{s}^{-1}$ | [28] |
| $K_{HCO_3^-}$ | Half-saturation constant for $HCO_3^-$ | $4.08 \times 10^{-4}$ | $\text{mol kg}^{-1}$ | [28] |
| $R_{z\_max}$ | Max respiratory rates of zooxanthellae | $1.6 \times 10^{-3}$ | $\text{mol m}^{-3} \text{s}^{-1}$ | Assumed<sup>a</sup> |
| $K_{CH_2O}$ | Half-saturation constant for $CH_2O$ | 0.1 | $\text{mol kg}^{-1}$ | Assumed<sup>a</sup> |
| $R_{c\_max}$ | Max respiratory rates of coral polyps | $0.4 \times 10^{-3}$ | $\text{mol m}^{-3} \text{s}^{-1}$ | Assumed<sup>a</sup> |
| $k_p$ | Precipitation rate constant | $1.1 \times 10^{-8}$ | $\text{mol m}^{-2} \text{s}^{-1}$ | [28] |
| $k_d$ | Dissolution rate constant | $2.7 \times 10^{-7}$ | $\text{mol m}^{-2} \text{s}^{-1}$ | [28] |
| $n_p$ | Reaction order for precipitation | 1.63 | / | [28] |
| $n_d$ | Reaction order for dissolution | 2.5 | / | [28] |
| $T_l$ | Temperature for lowest photosynthetic rate | 38.2 | $^{\circ}\text{C}$ | [44] |
| $T_h$ | Temperature for highest photosynthetic rate | 28 | $^{\circ}\text{C}$ | Assumed<sup>a</sup> |
| $n_T$ | Dimensionless parameter for sensitivity | 1.76 | / | [44] |
| $m$ | Algae distribution regulation parameter | 90 | $^{\circ}$ | [34] |
| $D_k$ | Mass diffusion coefficient of substance $k$ | | $\text{m}^2 \text{s}^{-1}$ | |
| $D_{DO}$ | In seawater | $2 \times 10^{-9}$ | $\text{m}^2 \text{s}^{-1}$ | [35] |
| | In coral | $1 \times 10^{-9}$ | $\text{m}^2 \text{s}^{-1}$ | [35] |
| $D_{DIC}$ | In seawater | $0.81 \times 10^{-9}$ | $\text{m}^2 \text{s}^{-1}$ | [47] |
| | In coral | $0.41 \times 10^{-9}$ | $\text{m}^2 \text{s}^{-1}$ | [47] |
| $D_{TA}$ | In seawater | $0.81 \times 10^{-9}$ | $\text{m}^2 \text{s}^{-1}$ | [47] |
| | In coral | $0.41 \times 10^{-9}$ | $\text{m}^2 \text{s}^{-1}$ | [47] |
| $D_{CH_2O}$ | In coral | $0.69 \times 10^{-9}$ | $\text{m}^2 \text{s}^{-1}$ | [48] |
| $\Delta G$ | Free energy released by ATP hydrolysis | $-3.05 \times 10^4$ | $\text{J mol}^{-1}$ | [28] |
| $\eta$ | Energy conversion efficiency | 0.3 | | [28] |
| $R_g$ | Gas constant | 8.314 | $\text{J mol}^{-1} \text{K}^{-1}$ | / |
| $k_{pp}$ | Conductivity through the paracellular pathway | $3 \times 10^{-5}$ | $\text{m s}^{-1}$ | [28] |

<sup>a</sup> These parameters were assumed to create a specific simulation system
<sup>b</sup> These parameters were roughly adjusted to match the experimental values of DO concentration and pH inside the coral [21, 49]

**Table A2.** The parameters with dynamically changing values Symbol Description Unit Ref.

| Symbol | Description | Unit | Ref. |
| --- | --- | --- | --- |
| $v_s$ | Coral surface velocity | $\text{m s}^{-1}$ | / |
| $\alpha$ | Angle | $^{\circ}$ | / |
| $\mathbf{u}$ | Fluid velocity vector | $\text{m s}^{-1}$ | / |
| $t$ | Time | s | / |
| $p$ | Fluid pressure | Pa | / |
| $\boldsymbol{\tau}$ | Stress tensor | $\text{N m}^{-2}$ | / |
| $c_k$ | Concentration of substance $k$ | $\text{mol kg}^{-1}$ | / |
| $f_{CO_2}$ | Fugacity of carbon dioxide in the atmosphere | Pa | [43] |
| $K_0$ | Equilibrium constants | $\text{mol kg}^{-1}$ | [43] |
| $K_1$ | Equilibrium constants | $\text{mol kg}^{-1}$ | [43] |
| $K_2$ | Equilibrium constants | $\text{mol kg}^{-1}$ | [43] |
| $K_w$ | Equilibrium constants | $\text{mol}^2 \cdot \text{kg}^{-1}$ | [43] |
| $E_d$ | Photon flux density reaching the coral surface | $\text{mol} \cdot \text{m}^{-2} \cdot \text{s}^{-1}$ | / |
| $\gamma$ | Solar altitude | ° | / |
| $T$ | Temperature | °C | / |
| $P$ | Photosynthetic rate of symbiotic algae | $\text{mol m}^{-3} \text{s}^{-1}$ | / |
| $k_T$ | Temperature parameters for photosynthesis | / | / |
| $k_E$ | Light intensity parameters for photosynthesis | / | / |
| $R_z$ | Respiratory rates of zooxanthellae | $\text{mol m}^{-3} \text{s}^{-1}$ | / |
| $k_z$ | Zooxanthellae concentration parameters for respiration | / | / |
| $R_c$ | Respiratory rates of coral polyps | $\text{mol m}^{-3} \text{s}^{-1}$ | / |
| $G$ | Calcification rate | $\text{mol m}^{-2} \text{s}^{-1}$ | / |
| $\Omega$ | Aragonite saturation state | / | / |
| $d$ | Distance from the coral polyps-calcified fluid interface | mm | / |
| $K_a$ | Solubility product for aragonite | $\text{mol}^2 \cdot \text{kg}^{-6}$ | [46] |
| $j_k$ | Diffusive flux of substance $k$ | $\text{mol m}^{-2} \text{s}^{-1}$ | / |
| $S_k$ | Source term of substance $k$ | $\text{mol m}^{-3} \text{s}^{-1}$ | / |
| $Q_R$ | Energy produced by respiration | $\text{J m}^{-3} \text{s}^{-1}$ | / |
| $Q_R^*$ | Average energy on the polyps-calcifying fluid interface | $\text{J m}^{-2} \text{s}^{-1}$ | / |
| $F_{TA}$ | TA flux from coral polyps to the calcifying fluid | $\text{mol m}^{-2} \text{s}^{-1}$ | / |
| $E_{H^+}$ | Energy required for $H^+$ transportation | $\text{J mol}^{-1}$ | / |
| $F_{TA\_pp}$ | TA flux through paracellular pathway | $\text{mol m}^{-2} \text{s}^{-1}$ | / |
| $F_{DIC\_pp}$ | DIC flux through paracellular pathway | $\text{mol m}^{-1} \text{s}^{-1}$ | / |
| $R_k$ | Mass transfer resistance of substance $k$ | $\text{s m}^{-1}$ | / |
| $\bar{c}_s$ | Average concentration on the coral surface | $\text{mol kg}^{-1}$ | / |
| $c_b$ | Bulk seawater concentration | $\text{mol kg}^{-1}$ | / |
| $\bar{F}_k$ | Surface average flux of substance $k$ ($\text{mol/m}^2 \cdot \text{s}$). | $\text{mol m}^{-2} \text{s}^{-1}$ | / |

